# A Broadly Neutralizing Macaque Monoclonal Antibody Against the HIV-1 V3-Glycan Patch

**DOI:** 10.1101/2020.08.10.244582

**Authors:** Zijun Wang, Christopher O. Barnes, Rajeev Gautam, Julio C. C. Lorenzi, Christian T. Mayer, Thiago Y. Oliveira, Victor Ramos, Melissa Cipolla, Kristie M. Gordon, Harry B. Gristick, Anthony P. West, Yoshiaki Nishimura, Henna Raina, Michael S. Seaman, Anna Gazumyan, Malcolm A. Martin, Pamela J. Bjorkman, Amelia Escolano, Michel C. Nussenzweig

**Author notes:** Equal contribution (Zijun Wang, Christopher O. Barnes, Rajeev Gautam). Address correspondence to Dr. Amelia Escolano.

## Abstract

A small fraction of HIV-1 infected humans develop potent broadly neutralizing antibodies (bNAbs) against HIV-1 that can protect macaques from infection with simian immunodeficiency HIV chimeric virus (SHIV). Similarly, a small number of macaques infected with SHIVs also develop broadly neutralizing serologic activity, but less is known about the nature of these simian antibodies. Here we report on a monoclonal antibody, Ab1485, isolated from a macaque infected with SHIV_AD8_ that developed broadly neutralizing serologic activity mapping to the V3-glycan region of HIV-1 Env. Ab1485 neutralizes 38.1 % of HIV-1 isolates in a panel of 42 pseudoviruses with a geometric mean IC_50_ of 0.055 μg/ml and SHIV_AD8_ with an IC_50_ of 0.028 μg/ml. Ab1485 binds to the V3-glycan epitope in a glycan-dependent manner as determined by ELISA and neutralization assays with JRCSF mutant viruses. A 3.5 Å cryo-electron microscopy structure of Ab1485 in complex with a native-like SOSIP Env trimer showed conserved contacts with the N332_gp120_ glycan and gp120 GDIR peptide motif, but in a distinct Env-binding orientation relative to human V3/N332_gp120_ glycan-targeting bNAbs. Finally, intravenous infusion of Ab1485 protected macaques from a high dose intrarectal challenge with SHIV_AD8_. We conclude that macaques can develop bNAbs against the V3-glycan patch that resemble human V3-glycan bNAbs.

**Significance statement:** Rhesus macaques infected with SHIV are an important model for evaluating HIV-1 prevention and therapy strategies and can also be used to evaluate humoral immune responses to candidate HIV-1 vaccines, but whether macaques produce human-like bNAbs has not been evaluated. Like HIV-1 infected humans, 10-20% of the SHIV_AD8_ challenged macaques develop low levels of neutralizing antibodies, and only one macaque has developed broad and potent serologic neutralizing activity. We have examined the antibody response of this macaque (CE8J) and we report on the cloning and molecular characterization of a bNAb produced in this elite neutralizing non-human primate, its structure bound to an HIV-1 Env trimer, and the implications for development of vaccines targeting the V3-glycan patch of Env.

## Introduction

Over the last decade, characterization of monoclonal antibodies from HIV-1-infected individuals with broad and potent serologic activity against the virus revealed that bNAbs are unusual in that nearly all carry large numbers of somatic mutations. Their target epitopes are also unusual because many combine host-derived glycans with protein components of the HIV-1 envelope spike protein (Env). Longitudinal cohort and structural studies demonstrated that bNAb somatic mutations arise in part to accommodate the host-derived glycans that shield Env (1–7), a process that requires multiple rounds of somatic mutation and selection driven by viral escape from immune pressure (2–5, 7–9).

The observation that bNAbs arise during natural infection in humans (10–17), and that they can block SHIV infection in macaques (18–28), suggests that a vaccine that elicits such antibodies would be protective. However, with the exception of genetically engineered mice (29–31), all such efforts have produced only sporadic or less than optimal results with little or no protective activity against heterologous viral strains(32, 33). More importantly, it remains unclear which animal model is most relevant to test candidate vaccines.

Non-human primates infected with SHIV_AD8_ develop high levels of prolonged viremia that leads to destruction of their CD4^+^ T cell compartment and an AIDS-like disease including immunodeficiency and infection with pneumocystis pneumonia and other opportunistic pathogens (34–37). Like HIV-1 infected humans, 10-20% of the SHIV_AD8_ challenged macaques develop low levels of neutralizing antibodies to a small number of heterologous strains, and only one has developed broad and potent serologic neutralizing activity (38). The serum from this macaque (CE8J) exhibited potent cross-clade anti-HIV-1 neutralizing activity similar to that observed for HIV-1-infected human elite neutralizers. This broadly neutralizing activity persisted throughout the 2-year infection of CE8J. This monkey ultimately succumbed to immunodeficiency and was euthanized 117 weeks post-infection. Plasma mapping studies revealed that neutralizing activity of CE8J macaque exclusively targeted the glycan patch associated with the variable 3 (V3) loop on HIV-1 Env (38).

Here we report on the cloning and molecular characterization of a bNAb produced in this elite neutralizing non-human primate, its structure bound to an HIV-1 Env trimer, and the implications for development of vaccines targeting the V3-glycan patch.

## Results

### Isolation of single Env-specific B cells from a SHIV_AD8_ infected macaque

To isolate bNAbs from macaque CE8J, we purified germinal center (GC) B cells that bound to YU2 gp140 fold-on trimer (YU2) and BG505 SOSIP.664 trimer (BG505), but not to a control antigen, from lymph nodes collected at the time of necropsy, 115 weeks post SHIV_AD8_ infection (Figure 1A and Figure S1A).

**Figure 1.**
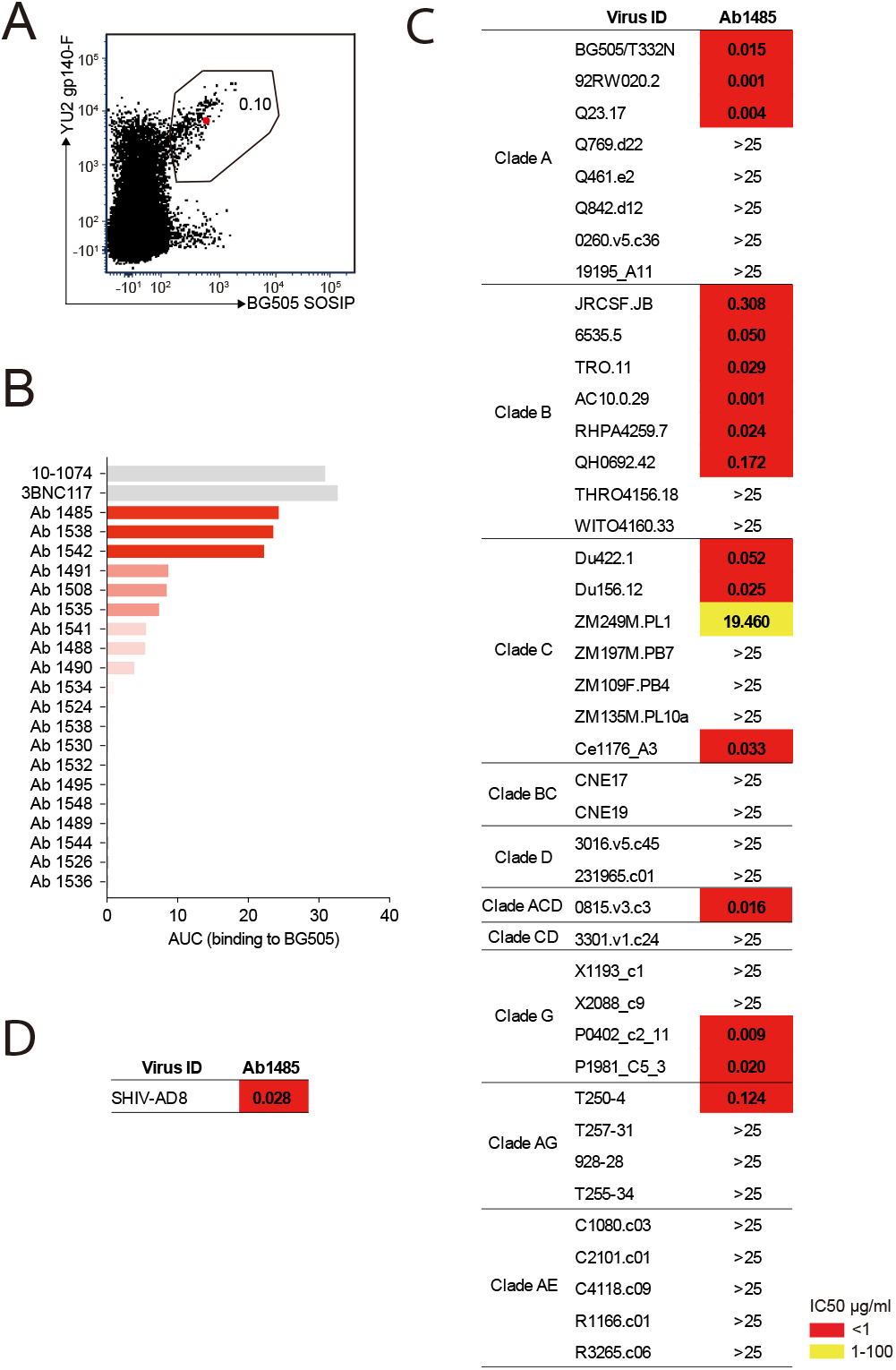
Broadly neutralizing antibody isolated from a SHIV_AD8_ infected rhesus macaque. **A**, FACS plot showing germinal center B cells that bind to YU2 gp140-F and BG505 SOSIP from a lymph node collected from macaque CE8J at week 115 after SHIV_AD8_ infection. The gate shows the sorting window. The B cell carrying the isolated bNAb (Ab1485) is highlighted in red. **B**, Graph shows the binding of several monoclonal antibodies isolated from macaque CE8J to BG505 SOSIP. Data is shown as area under the ELISA curve (AUC). **C and D**, Table shows the neutralization activity of Ab1485 determined in TZM-bl assays against a panel of 42 multi clade tier 1B and tier 2 pseudoviruses (C) and replication competent SHIVAD8 (D).

Immunoglobulin heavy chain (IgH) and light chains Lambda (IgL) and Kappa (IgK)-encoding mRNAs were amplified from the isolated Env-specific B cells by PCR using a set of primers specifically designed to amplify a diverse set of macaque genes (39). Paired IgH and IgL/IgK sequences were obtained for 90 antibodies. Among the 90 antibodies, we found two expanded B cell clones (Figure S1B). Sequence analysis revealed that IgH, IgL and IgK genes were somatically hypermutated (averages of 11.8, 8.5 and 4.1 average nucleotide mutations, respectively) (Figure S1C). The average length of the complementarity determining region 3 of the heavy chain (CDRH3) was 15.3 amino acids, and 30 of the antibodies had CDRH3s of 18 or more amino acids (Figure S1D).

### Ab1485 isolated from macaque CE8J is potently neutralizing

We produced 67 of the 90 monoclonal antibodies and tested them for binding to BG505 by ELISA. Several antibodies showed detectable binding to BG505 with Ab1485 showing the highest-level binding (Figure 1B). The three best binders to BG505 were evaluated for neutralizing activity in TZM-bl assays (40) against a screening panel of 7 HIV-1 pseudoviruses. Only Ab1485 showed potent and broad activity against this panel (Figure S1E). To further evaluate the neutralization potency and breadth of Ab1485, we tested it in TZM-bl assays against a panel of 42 pseudoviruses covering 9 different HIV-1 clades (Figure 1C). Ab1485 neutralized 16 of the isolates in the 42-virus panel with a mean IC_50_ of 0.055 μg/mL (Figure 1C), and it was also a potent neutralizer of a replication-competent SHIV (SHIV_AD8_, IC_50_ = 0.028 μg/mL, Figure 1D). We conclude that Ab1485 is a potent neutralizer with limited breadth compared to some of the human bNAbs reported to date (41).

### Mapping the Ab1485 epitope on HIV-1 Env

Ab1485 combines the germline V gene segments VH4_2M and VL124_30 and is highly mutated (33 and 25 nucleotide mutations in the VH and VL genes respectively). It has a 20-residue CDRH3 (Figure 2A).

**Figure 2.**
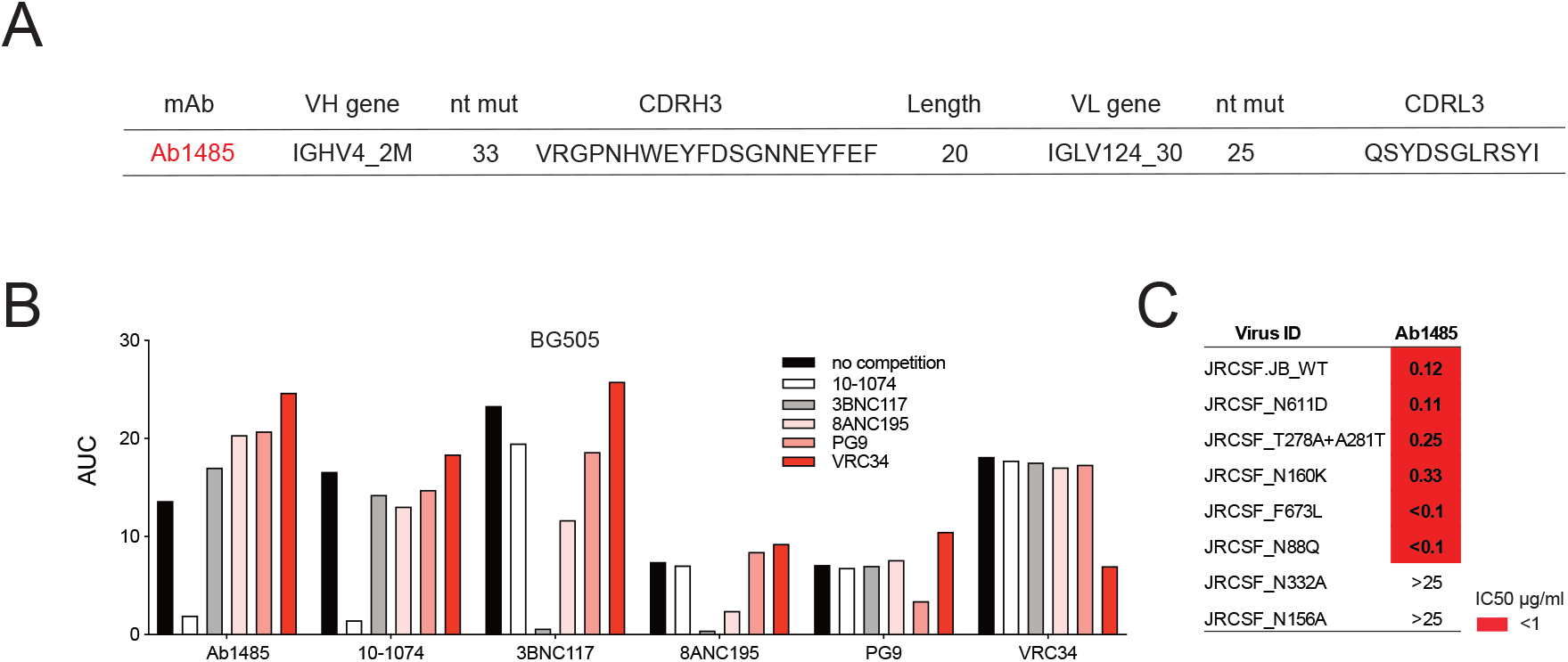
Mapping of Ab1485 binding to Env. **A**, Description of Ab1485. **B**, ELISA binding of Ab1485 to BG505 in competition with antibodies that target the V3-glycan epitope (10-1074), the CD4 binding site (3BNC117), the gp120-gp41 interface (8ANC195), the apex (PG9) or the fusion peptide (VRC34) and in the absence of a competing antibody. **C**, Table shows the neutralization activity of Ab1485 determined in TZM-bl assays against a JRCSF pseudovirus and a series of JRCSF mutants that affect the binding of human bNAbs to the interface (N611D), the CD4 binding site (T278A+ A281T), the apex (N160K), the MPER (F673L) and the fusion peptide (N88Q).

To characterize the target epitope of Ab1485, we performed competition ELISAs using the bNAbs 10-1074, 3BNC117, 8ANC195, PG9, VRC34 that target the V3-glycan patch, the CD4 binding site, the gp120-gp41 interface, the apex and the fusion peptide of Env, respectively (42–45). Binding of these antibodies was self-inhibitory and in addition, 3BNC117 was partially inhibited by the gp120-gp41 interface bNAb 8ANC195 (43, 46, 47), and vice versa. The binding of Ab1485 to BG505 was inhibited by the V3-glycan bNAb 10-1074 but not by any of the other human bNAbs (Figure 2B).

To further map the antibody target site, we performed neutralization assays using a series of HIV-1_JRCSF_ mutants (39). The neutralizing activity of Ab1485 was dependent on the potential N-linked glycosylation sites at N332_gp120_ and N156_gp120_, but unaffected by mutations that interfere with the binding of human bNAbs to the interface (N611D), the CD4 binding site (T278A+ A281T), the V1V2 apex (N160K), the MPER (F673L) or the fusion peptide (N88Q) (Figure. 2C). In conclusion, the competition ELISA and neutralization results suggested that Ab1485 targets the V3-glycan patch in Env.

### Cryo-EM structure of Ab1485 in complex with BG505

To elucidate the molecular details of Env recognition by Ab1485, we determined a 3.5 Å cryo-EM structure of Ab1485 in complex with the BG505 SOSIP.664 trimer and the gp120-gp41 interface targeting antibody, 8ANC195 (46, 47) (Figure 3A, Table S1 and Figure S2). The structure of the Ab1485-Env complex revealed recognition of an epitope focused on the N332_gp120_ glycan, the gp120 GDIR peptide motif, and V1 loop (Figure 3B). In common with human-derived V3/N332_gp120_ glycan-targeting bNAbs, Ab1485’s primary interaction was with the N332_gp120_ glycan (Man_6_GlcNAc_2_), which interfaces almost entirely with the CDRH3 loop (Figure 3C) (~400 Å^2^ antibody buried surface area (BSA)). In addition to the N332_gp120_ glycan, Ab1485 makes secondary contacts with the N156_gp120_ glycan (~275 Å^2^ antibody BSA), which frames Ab1485’s VH domain at the gp120 V3 epitope, rationalizing the observed loss in neutralization activity when tested against the JRCSF ΔN156_gp120_ glycan virus (Figure 2C) and consistent with faster dissociation kinetics observed in surface plasmon resonance (SPR) binding experiments that demonstrated a faster dissociation rate for a SOSIP that lacks the N156_gp120_ glycan (RC1) compared to a SOSIP that contains the N156_gp120_ potential N-linked glycosylation site (BG505) (Figure S3A-C).

**Figure 3.**
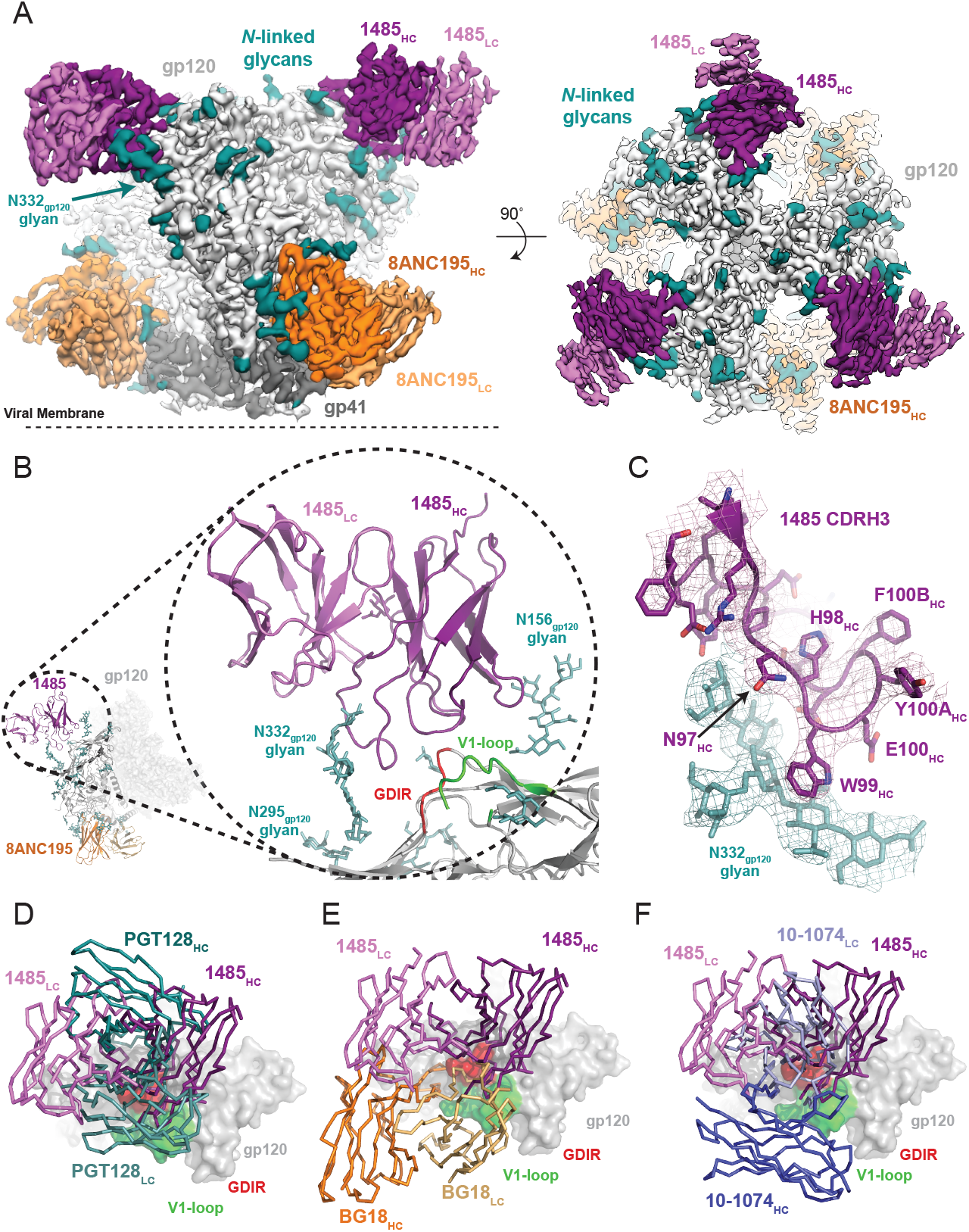
Cryo-EM reconstruction of the Ab1485-BG505 complex reveals a distinct Env binding orientation relative to human bNAbs. (A) Cryo-EM map of the BG505 SOSIP.664 trimer bound to three Ab1485 (purple shades) and three 8ANC195 (orange shades) Fabs. Densities for glycans are colored in dark teal. (B) Cartoon depiction of the modeled complex with a close-up view of the Ab1485 Fab - gp120 interface. Conserved regions of the V3-epitope are highlighted. (C) Cartoon and stick representation of the Ab1485 CDRH3 recognition of the N332gp120-glycan. Reconstructed cryo-EM map shown as a mesh, contoured at 3 sigma. (D-F) Comparison of Ab1485’s (purple) Env-binding orientation to (D) PGT128 (teal, PDB 5ACO), (E) BG18 (orange, PDB 6CH7), and (F) 10-1074 (blue, PDB 5T3Z).

Despite sharing a common interaction with the N332_gp120_ glycan, numerous studies have demonstrated that the V3-targeting bNAbs can adopt different binding orientations when targeting the N332_gp120_ glycan supersite(1, 48, 49). When compared to the poses of human-derived bNAbs (Figure 3D-F) and V3-targeting antibodies elicited in rabbits or rhesus macaques by vaccination or SHIV_BG505_ challenge (Figure S3D), Ab1485 adopts a Env-binding orientation in a manner most closely related to PGT128, which primarily uses its heavy chain to interact with the V3/N332_gp120_-glycan epitope (50) (Figure 3D-F). However, in contrast to PGT128, Ab1485 lacks any V-gene insertions or deletions and its orientation is rotated ~90° relative to PGT128, resulting in a unique Env-binding orientation that shifts interactions away from the N301_gp120_ N-glycan and towards the N156_gp120_ glycan, and moves the light chain outside of the underlying V3-epitope (Figure 3 and Figure 4A-C). Thus, interactions with the N-linked glycans and gp120 peptide components are almost exclusively mediated by the Ab1485 heavy chain (Figure 4A,B; 1255 Å^2^ vs 65 Å^2^ buried surface areas for HC and LC components of Ab1485 paratope, respectively). This observation suggests that Ab1485 may not be restricted by light chain pairing or require the consensus light chain sequence motifs commonly observed in human-derived V3/N332_gp120_-glycan targeting bNAbs.

**Figure 4.**
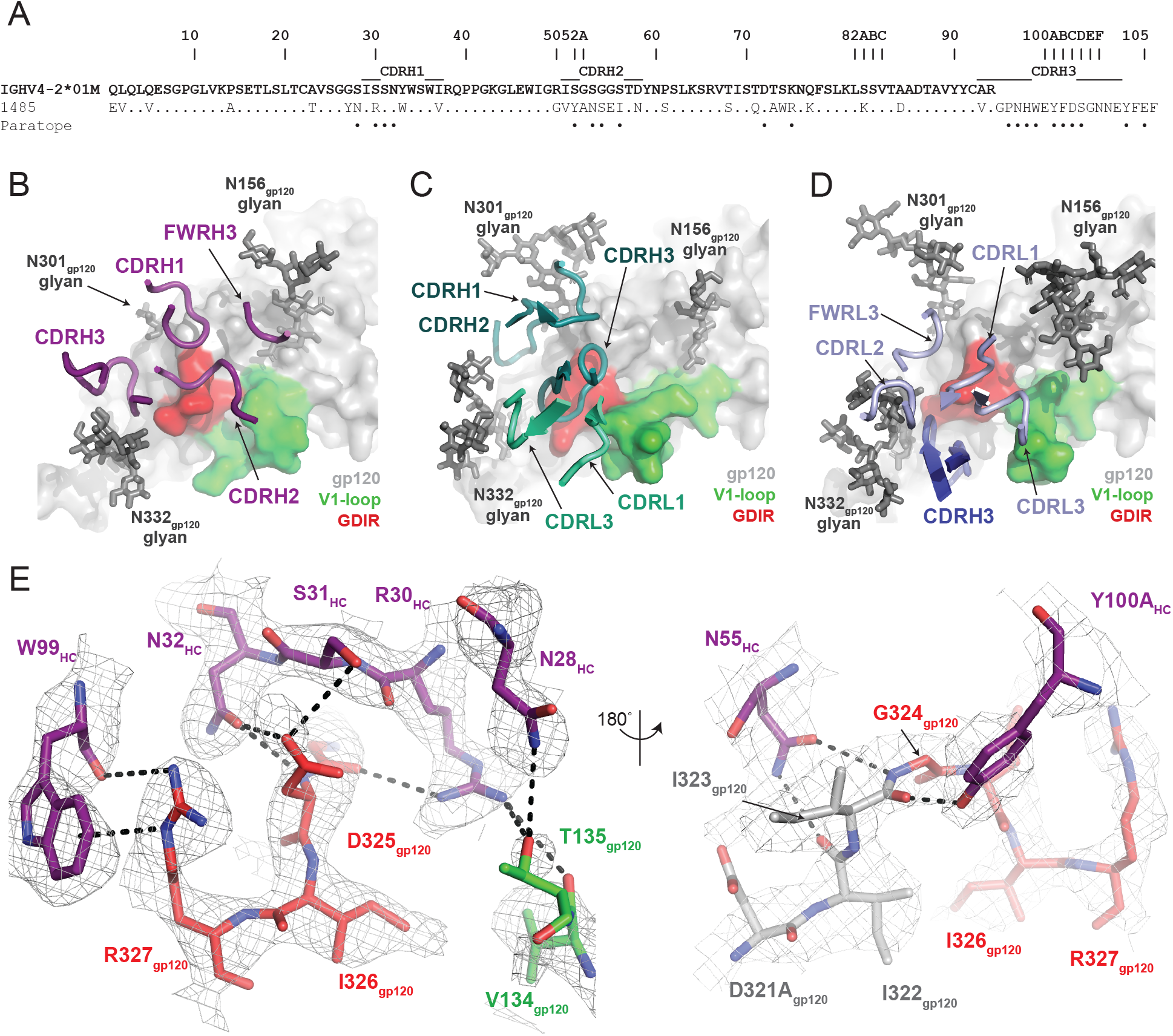
Molecular details of Ab1485-gp120 interactions. (A) Sequence alignment of mature Ab1485 heavy chain with germline VH4-2*01M. Paratope residues are highlighted. (B-D) Comparison of the paratope CDR loops and FWRs involved in epitope recognition for (B) Ab1485, (C) PGT128, and (D) 10-1074. (E) Stick representation of interactions between Ab1485 (purple) and either the GDIR peptide motif (red) or V1 loop (green). Potential H-bonds defined are shown as black dashes

Closer examination of the Ab1485 epitope revealed that all three heavy chain CDR loops contact the 32_4_GDIR_327_ gp120 peptide motif at the base of the V3-loop, which contrasts interactions by both heavy and light chain CDR loops observed in 10-1074/PGT121-like and BG18 bNAbs(51, 52) (Figure 4B,D). The primary molecular contacts are with CDRH1 residues R30_HC_, S31_HC_, and N32HC, which form potential hydrogen bonds with backbone and sidechain atoms from residues G324_gp120_ and D325_gp120_ (Figure 4E). Interestingly, these CDRH1 residues mimic interactions made by CDRL1/L3 residues in PGT121/101074-like bNAbs with Env, providing evidence for a convergent chemical mechanism of interactions with the gp120 GDIR motif, as previously shown for BG18 (52). Moreover, Ab1485 residue R30_HC_ forms secondary interactions with V1 loop residues V134_gp120_ and T135_gp120_, resembling similar arginine-gp120 V1-loop interactions observed in 10-1074, BG18, and PGT128(50–52).

A common interaction seen in the PGT121/101074-like, BG18, and PGT128 bNAbs involves the formation of a salt bridge between R327_gp120_ and an either a glutamate or aspartate in CDRH3 (42, 52). The Ab1485-BG505 structure reveals an alternate binding mode that involves the guanidinium moiety of R327_gp120_ forming cation-pi stacking interactions with W99_HC_ at the tip of the Ab1485’s CDRH3, while also participating in hydrogen bonding with the backbone carbonyl group (Figure 4E). While CDRH3 residue E100_HC_ could potentially adopt a conformation that would promote salt bridge formation with R327_gp120_, this residue is pointed outward and within H-bonding distance to the N332_gp120_-glycan in the Ab1485-BG505 complex. This observation suggests that salt-bridge formation between R327_gp120_ and an acidic CDRH3 residue found in V3-targeting bNAbs may not be as critical to targeting the V3/N332gp120-glycan epitope.

### Ab1485 protects from infection by SHIV_AD8_ in rhesus macaques

To determine whether Ab1485 can protect against SHIV_AD8_ infection in macaques, we expressed a fully macaque IgG, including Fc region substitutions that increase half-life through rescue by increased FcRn recycling (Ab1485-LS) (53). Ab1485 was not polyreactive, as shown by ELISAs against a series of antigens (Figure S4A) and Ab1485-LS had a half-life of 2.67 days in transgenic human FcRn mice (54) (Figure S4B). The protective efficacy of Ab1485-macaque-LS was assessed in rhesus macaques that received a single high dose of SHIV_AD8_ intrarectally (1000 TCID50) one day after a single intravenous infusion of Ab1485 at 10 mg/kg body weight (Figure 5A). Two historical control monkeys(55) receiving no mAb, rapidly became infected, generating peak levels of plasma viremia on day 14 post challenge. In contrast, the 4 macaques receiving Ab1485 remained uninfected throughout a 25-week observation period (Figure 5B). Neutralizing antibody titers persisted in the peripheral blood for at least 50 days after the virus challenge (Figure 5C). We conclude that Ab1485-macaque-LS protects macaques from high dose intrarectal SHIV_AD8_ infection.

**Figure 5.**
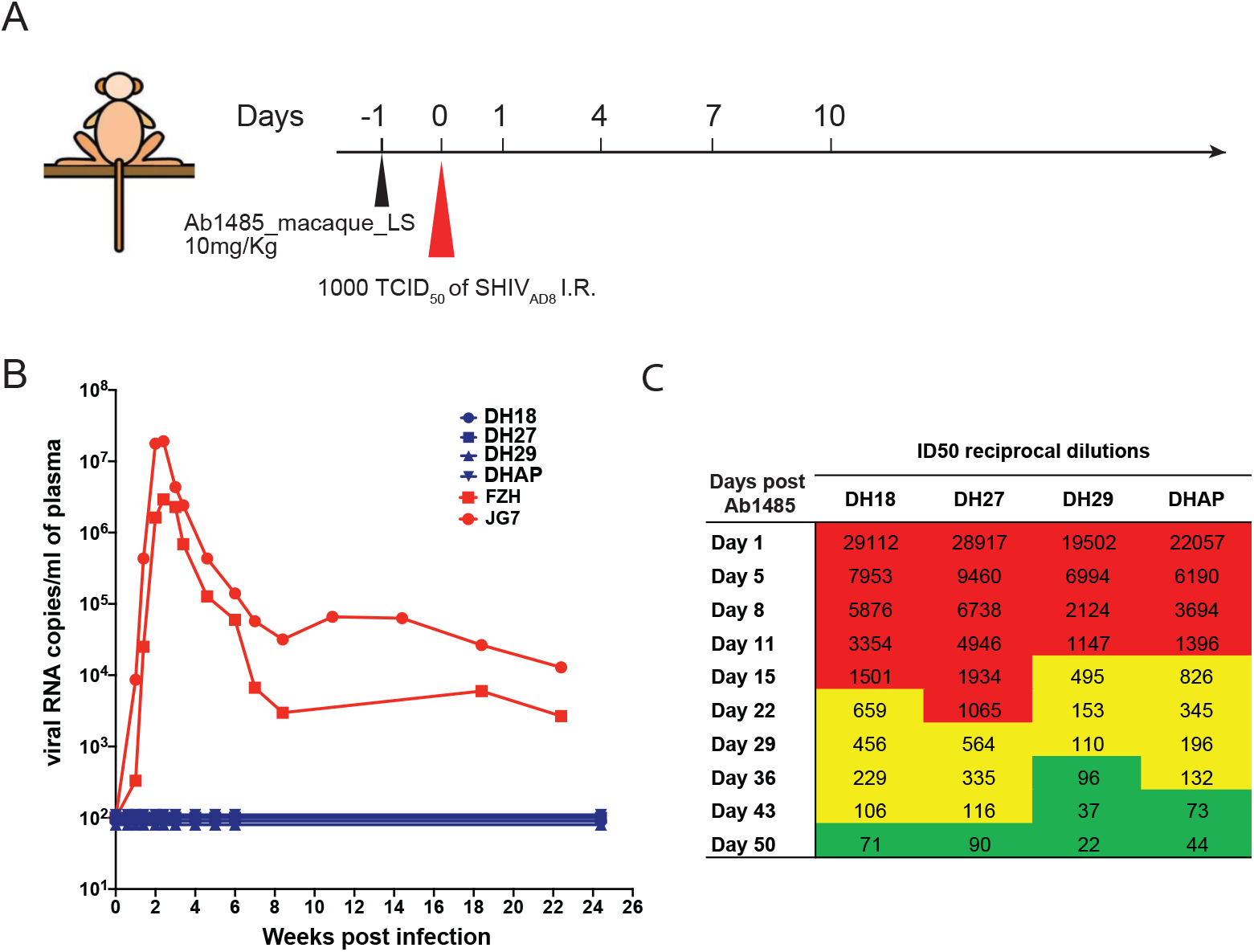
Ab1485 protects macaques against a high dose intrarectal challenge with SHIV_AD8_. **A**, Diagrammatic representation of the regimen used to assess the protective efficacy of Ab1485. Macaques were administered with Ab1485 at a dose of 10 mg kg^-1^ and challenged one day later with 1000 TCID_50_ of SHIV_AD8_ intrarectally (I.R.) **B**, Longitudinal analysis of plasma viral loads in two control macaques (FZH and JG7) receiving no Ab and four macaques (DH18, DH27, DH29 and DHAP) infused with Ab1485 24h prior to SHIV_AD8_ challenge. **C**, Serum neutralizing antibody titers in macaques receiving Ab1485. The IC_50_ titers are color coded: 1:21-99 in green; 1:100-999 in yellow and ≥ 1:1000 in red.

## Discussion

Indian-origin rhesus macaques infected with SHIV are an important model for evaluating HIV-1 prevention and therapy strategies. Macaques can also potentially be used to evaluate humoral immune responses to candidate HIV-1 vaccines, but whether macaques produce human-like bNAbs has not been evaluated. We have examined the antibody response of macaque CE8J who developed broad and potent serologic activity against HIV-1 many months after infection with SHIV_AD8_. Monoclonal Ab1485 obtained from single B cells purified on the basis of HIV-1 Env binding by cell sorting neutralized 16 of 42 of the HIV-1 strains tested. Through biochemical and structural analysis, we determined that antibody 1485 targets the V3/N332_gp120_-glycan epitope and does so in a manner that is similar to the human V3-targeting antibodies. Thus, rhesus macaques develop anti-HIV-1 V3-glycan patch bNAbs that are related to human bNAbs.

As many as 10-20% of HIV-1 infected humans develop antibodies that can neutralize a number of different HIV-1 strains, but only an elite few (1-2%) produce potent broadly neutralizing serologic activity(56). The elite humoral responders typically take 1-3 years to produce bNAbs, which is highly unusual for an antibody response. Single cell antibody cloning revealed that human bNAbs carry large numbers of somatic mutations that are required for their neutralizing activity(42, 43, 57–65). This observation led to the proposal that the development of bNAbs required a prolonged series of sequential interactions between the antibody-producing B lymphocytes and virus escape variants to drive antibody maturation(57, 66). Elegant prospective studies of virus and antibody evolution supported this concept (3–5, 7, 8), and sequential immunization experiments reproduced it in knock in mice (30). Monoclonal antibody Ab1485 resembles human bNAbs in many important respects including the high levels of somatic mutation suggesting a requirement for sequential interaction between B cells and the virus to drive bNAb evolution in macaques.

The glycan patch that surrounds the conserved GDIR peptide at the base of the V3 loop comprises an epitope that is frequently targeted by human bNAbs. Monoclonal antibody Ab1485 resembles PGT128 in that most of the interactions with the V3/N332_gp120_-glycan epitope are mediated by heavy chain CDR loops, including conserved interactions with the N332_gp120_ glycan and gp120 GDIR peptide motif. Recent studies in macaques showed that after priming with a designed immunogen that focuses the response to the V3-glycan epitope, antibodies with binding features of V3-glycan targeting bNAbs can be isolated(39). Whether these early antibodies can mature to develop broad neutralizing activity remains to be determined, but antibodies like Ab1485 may provide a blueprint for achieving such broad and potent activity.

Some of the predicted impediments for binding of germline forms of PGT121/10-1074-like or BG18-like antibodies are related to overcoming unfavorable interactions with the antibody light chain, particularly with the gp120 V1 loop. The unique orientation adopted by Ab1485, which positions the LC away from the V1 loop, may provide an easier path towards antibody maturation. Overall, structural analysis of 1485 provides: (i) evidence that effective bNAbs targeting the V3/N332_gp120_-glycan epitope are not restricted to PGT128, BG18 or 10-1074/PGT121-like binding mechanisms, (ii) critical structural insights towards immunogen design efforts to elicit neutralizing antibodies, including alternative binding modes that do not include light chain interactions, and (iii) evidence that SHIV_AD8_-infected macaques are capable of generating bNAbs that target the V3/N332_gp120_-glycan epitope in a manner similar to human-derived bNAbs, and therefore represent an excellent animal model for developing HIV-1 vaccines targeting this site.

Human bNAbs can protect humanized mice and macaques from HIV-1 and SHIV infections, respectively, when given prophylactically (18–28, 67, 68). They can also suppress infection for prolonged periods of time in mice and macaques(34, 69–71). When administered to Indian origin rhesus macaques during the acute SHIVAD8 infection they induced host CD8+ T cell-dependent immunity that can suppress infection for 2 to 3 years(70). However, prolonged administration of these human monoclonal antibodies to macaques has not been possible due to rapid development of macaque anti-human antibody responses. The fully-native macaque Ab1485 will facilitate such therapies and reservoir reduction experiments that require prolonged bNAb administration to macaques.

Finally, the observation that Indian origin rhesus macaques develop V3-glycan patch bNAbs that are closely related to human bNAbs is strongly supportive of the use of this model organism to test HIV-1 candidate vaccines that target this epitope.

## Materials and Methods

### Flow cytometry and single B cell sorting

Frozen lymph node (LN) cell suspensions collected from macaque CE8J at 117 weeks post-infection were thawed and incubated in FACS buffer (1 X Phosphate-buffered saline (PBS), 2% calf serum, 1 mM EDTA.) with the following anti human antibodies: anti CD3-APC-eFluor 780 (Invitrogen, 47-0037-41), anti CD14-APC-eFluor 780 (Invitrogen, 47-0149-42), anti CD16-APC-eFluor 780 (Invitrogen, 47-0168-41), anti CD8-APC-eFluor 780 (Invitrogen, 47-0086-42), anti CD20-PE-Cy7 (BD biosciences, 335793), and anti CD38 FITC (Stem Cell, 60131FI) and the Zombie NIR™ Fixable Viability Kit (Biolegend, 77184). Avi-tagged and biotinylated BG505 SOSIP, YU2-gp140 fold-on, and hepatitis B surface antigen (HBs Ag) (Protein Specialists, hbs-875) conjugated to streptavidin Alexa Fluor 647 (Biolegend, 405237), streptavidin PE (BD biosciences, 554061) and streptavidin BV711(BD biosciences, 563262) respectively were added to the antibody mixture and incubated with the LN cells for 30 min. Single CD3^-^CD8^-^CD14^-^ CD16^-^ CD20^+^CD38^+^ BG505 SOSIP^+^YU2-gp140^+^ B cells were sorted into individual wells of a 96-well plates containing 5 μl of a lysis buffer (Qiagen, 1031576) per well using a FACS Aria III (Becton Dickinson). The sorted cells were stored at −80°C or immediately used for subsequent RNA purification(39, 72).

### Single-B-cell antibody cloning

Single cell RNA was purified from individual B cells using magnetic beads (Beckman Counter, RNAClean XP, A63987) and used for cDNA synthesis by reverse transcription (SuperScript III Reverse Transcriptase, Invitrogen, 18080-044, 10,000 U). cDNA was stored at −20 °C or used for subsequent amplification of the variable IgH, IgL and IgK genes by nested PCR and SANGER sequencing using the primers and protocol previously described (39).

IgH, IgL and IgK V(D)J genes were cloned into expression vectors containing the human IgG1, IgL or IgK constant region using sequence and ligation independent cloning (SLIC) as previously described (39, 73). Ab1485-macaque-LS was cloned using pre designed gene fragments (IDT).

### Antibody production

IgGs were expressed by transient transfection in HEK293-6E cells and purified from cell supernatants using protein A or G (GE Healthcare) as previously described (39, 72).

### ELISA

ELISAs using BG505 SOSIP directly coated on a 96-well plate (Life Sciences, #9018) were performed as previously described (39). Briefly, high affinity 96-well flat bottom plates were coated with the SOSIP at 2 μg/ml and incubated overnight at 4°C. The plates were washed 3 times with PBS-0.05% Tween 20 and blocked with 2% of milk for 1 hour at RT. After blocking, monoclonal antibodies were added to the plate at 3-fold serial dilutions starting at 10 μg/ml and incubated for 2h at RT. Binding was developed with a horseradish peroxidase (HRP)-conjugated anti-human IgG secondary antibody (Jackson ImmunoResearch, 109-035-088) and using ABTS as the HRP substrate.

For competition ELISA, 96-well flat bottom plates were first coated with streptavidin (2 μg/ml) at 37°C for 1 h, then washed and blocked with 2%milk and incubated with biotinylated BG505 SOSIP (2 μg/ml) at 37°C for 1 h. Competing bNAb Fabs (10-1074, 3BNC117, 8ANC195, VRC34 and PG9) were added at 10 μg/ml to the plates for 1 h at 37C. After wash, serially diluted mAbs were added and incubated at 37°C for 2 h. The binding was detected by an HRP-conjugated anti-human IgG (Fc) CH2 Domain antibody (Bio-Rad MCA647P) used at a 1:5,000 dilution at RT for 1 h and developed as described above.

### Polyreactivity assay

ELISAs to determine antibody binding to LPS, KLH, single stranded DNA, double stranded DNA and insulin were previously described in (74). ED38 (75, 76) and mG053(77) antibodies were used as positive and negative controls.

### Antibody pharmacokinetic analysis

Female B6. Cg-Fcgrt^tm1Dcr^ Tg(CAG-FCGRT)276Dcr/DcrJ mice (FcRn -/- hFcRn) (The Jackson Laboratory, #004919) aged 7-8 weeks were intravenously injected (Retro-orbital vein) with 0.5 mg of purified Ab1485-macaque-LS in PBS. Total serum concentrations of human IgG were determined by ELISA as previously described with minor modifications (78). In brief, high-binding ELISA plates (Corning) were coated with Goat Anti-Human IgG (Jackson ImmunoResearch #109-005-098) at a concentration of 2.5 μg/ml overnight at RT. Subsequently, wells were blocked with blocking buffer (2% BSA (SIGMA), 1 mM EDTA (Thermo Fisher), and 0.1% Tween 20 (Thermo Fisher) in PBS). A standard curve was prepared using human IgG1 kappa purified from myeloma plasma (Sigma-Aldrich). Serial dilutions of the IgG standard (in duplicates) and serum samples in PBS were incubated for 60 min at 37C, followed by HRP-conjugated anti-human IgG (Jackson ImmunoResearch #109-035-008) diluted 1: 5,000 in blocking buffer for 60 min at RT. Following the addition of TMB (Thermo Fisher #34021) for 8 minutes and stop solution, optical density at 450 nm was determined using a microplate reader (BMG Labtech). Plates were washed with 0.05% Tween 20 in PBS between each step. The elimination half-life was calculated using pharmacokinetics parameters estimated by performing a non-compartmental analysis (NCA) using WinNonlin 6.3 (Certara Software).

### *In vitro* neutralization assays

The neutralization activity of monoclonal antibodies was assessed using TZM-bl cells as described previously(79).

### Protein expression and purification for structural studies

Ab1485 and 8ANC195 Fabs were recombinantly expressed by transiently transfecting Expi293F cells (Invitrogen) with vectors encoding antibody light chain and C-terminal hexahistidine-tagged heavy chain genes. Secreted Fabs were purified from cell supernatants harvested four days post-transfection using Ni^2+^-NTA affinity chromatography (GE Healthcare), followed by size exclusion chromatography (SEC) with a Superdex 16/60 column (GE Healthcare). Purified Fabs were concentrated and stored at 4 °C in storage buffer (20 mM Tris pH 8.0, 120 mM NaCl, 0.02% sodium azide).

A gene encoding soluble BG505 SOSIP.664 v4.1 gp140 trimer (80) was stably expressed in Chinese hamster ovary cells (kind gift of John Moore, Weill Cornell Medical College) as described (81). Secreted Env trimers were isolated using PGT145 immunoaffinity chromatography by covalently coupling PGT145 IgG monomer to an activated-NHS Sepharose column (GE Healthcare) as previously described (81). Properly folded trimers were eluted with 3M MgCl_2_ and immediately dialyzed into storage buffer before being subjected to multiple size exclusion chromatography runs with a Superdex200 16/60 column followed by a Superose6 10/300 column (GE Healthcare). Peak fractions verified to be BG505 SOSIP.664 trimers were stored as individual fractions at 4°C in storage buffer.

### Cryo-EM sample preparation

Complexes of Ab1485-BG505-8ANC195 were assembled by incubating purified Fabs with BG505 trimers at a 3:1 Fab:gp140 protomer ratio overnight at room temperature. Complexes were purified from excess Fab by size exclusion chromatography using a Superose-6 10/300 column (GE Healthcare). Peak fractions corresponding to the Ab1485-BG505-8ANC195 complex were concentrated to 1.5 mg/mL in 20 mM Tris pH 8.0, 100 mM NaCl and deposited onto a 400 mesh, 1.2/1.3 Quantifoil grid (Electron Microscopy Sciences) that had been freshly glow-discharged for 45 s at 20 mA using a PELCO easiGLOW (Ted Pella). Samples were vitrified in 100% liquid ethane using a Mark IV Virtobot (Thermo Fisher) after blotting for 3s with Whatman No. 1 filter paper at 22°C and 100% humidity.

### Cryo-EM data collection, processing, and model refinement

Movies of the Ab1485-BG505-8ANC195 complex were collected on a Talos Arctica transmission electron microscope (Thermo Fisher) operating at 200 kV using SerialEM automated data collection software (82) and equipped with a Gatan K3 Summit direct electron detector. Movies were obtained in counting mode at a nominal magnification of 45,000x (super-resolution 0.435 Å/pixel) using a defocus range of −1.5 to −3.0 μm, with a 3.6 s exposure time at a rate of 13.4 e^-^/pix/s, which resulted in a total dose of 60 e^-^/Å^2^ over 40 frames.

Cryo-EM data processing was performed as previously described (83). Briefly, movie frame alignment was carried out with MotionCorr2 (84) with dose weighting, followed by CTF estimation in GCTF (85). After manual curation of micrographs, reference-free particle picking was conducted using Laplacian-of-Gaussian filtering in RELION-3.0. Extracted particles were binned x4 (3.47 Å/pixel), and subjected to reference-free 2D classification with a 220 Å circular mask. Class averages that represented different views of the Fab bound Env-trimer and displayed secondary structural elements were selected (~641,000 particles) and an ab initio model was generated using cryoSPARC v2.12 (86).

The generated volume was used as an initial model for iterative rounds of 3D classification (C1 symmetry, k=6) in RELION 3.0. Particles corresponding to 3D class averages that displayed the highest resolution features were re-extracted unbinned (0.869 Å/pixel) and homogenously 3D-refined with a soft mask in which Fab constant domains were masked out, resulting in an estimated resolution of 4.1 Å according to gold-standard FSC (87). To further improve the resolution, particles were further 3D classified (k=6, tau_fudge=10), polished, and CTF refined. The final particle stack of ~404,000 particles refined to a final estimated resolution of 3.54 Å (C1 symmetry) according to gold-standard FSC.

To generate initial coordinates, reference models (gp120-gp41, PDB: 6UDJ; 8ANC195 Fab, 4PNM) were docked into the final reconstructed density using UCSF Chimera v1.13 (88). For the Ab1485, initial coordinates were generated by docking a 10-1074 Fab model (PDB 4FQQ), which had been altered by removing Fab CDR loops, into the cryo-EM density at the V3/N332-glycan epitope. Prior to initial refinement, 10-1074 VH-VL sequences were mutated to match Ab1485 and manually refined into density in Coot(89). The full model was then refined into the cryo-EM maps using one round of rigid body refinement, morphing, and simulated annealing followed by several rounds of B-factor refinement in Phenix(90). Models were manually built following iterative rounds of real-space and B-factor refinement in Coot and Phenix with secondary structure restraints. Modeling of glycans was achieved by interpreting cryo-EM density at possible N-linked glycosylation sites in Coot. Validation of model coordinates was performed using MolProbity(91) and figures were rendered using UCSF Chimera or PyMOL (Version 1.5.0.4 Schrodinger, LLC). Buried surface areas and potential hydrogen bonds were determined as previously described (83).

### Surface Plasmon Resonance

SPR experiments were performed using a Biacore T200 instrument (GE Healthcare). RC1 SOSIP (39), RC1 glycan-KO ^324^GAIA^327^SOSIP (39) and BG505 SOSIP (39) (were immobilized on a CM5 chip by primary amine chemistry (Biacore manual). Flow cell 1 was kept empty and reserved as a negative control. A concentration series of 1485 Fab (3-fold dilutions from a top concentration of 100 nM) was injected at 30 μl/min for 60s followed by a dissociation phase of 300s. Binding reactions were allowed to reach equilibrium and *K_D_* values were calculated from the ratio of association and dissociation constants (*K*_D_ = *k*_d_/*k*_a_), which were derived using a 1:1 binding model that was globally fit to all curves in a data set. Flow cells were regenerated with 10 mM glycine pH 3.0 at a flow rate of 90 μl/min for 30s.

### Virus Challenge

A single dose (10 mg/kg body weight) of Ab1485 was infused intravenously into four Indian rhesus macaques (DH18, DH27, DH29 and DHAP). 24 h following Ab infusion, these animals were inoculated intrarectally with a high dose (1000 TCID_50_) challenge of SHIV_AD8_. Two control monkeys (FZH and JG7), receiving no Ab, were reported in a previous study(55). A pediatric speculum was used to gently open the rectum and a 1 ml suspension of virus in a tuberculin syringe was slowly infused into rectal cavity. Blood was drawn regularly to monitor viral infection and serum neutralizing activity. All animal procedures and experiments were performed according to protocols approved by the Institutional Animal Care and Use Committee of NIAID, NIH.

### Analysis

MacVector v.17.0.2 was used for sequence analysis. Flow cytometry data were processed using FlowJo v.10.6.1and FCS EXPRESS. GraphPad Prism 7 was used for data analysis. Immunoglobulin gene sequence AB1 files were converted to FASTQ format using SeqIO from Biopython (92). In the quality control step, non-determined and low-quality nucleotides were trimmed from both 5’ and 3’ ends of the sequence present in the FASTQ files using cutadapt v.2.3 software. IgBlast was used to identify immunoglobulin V(D)J genes and consequently the junction region, which was further used to define the Ig clones by Change-O toolkit v.0.4.5(93). Clones were defined by calculating and normalizing the hamming distance of the junction region and comparing it to a pre-defined threshold of 0.15.

## Author contributions

Z.W., C.O.B., R.G., M.A.M., P.J.B., M.C.N. and A.E. designed the research. Z.W., C.O.B., R.G., J.C.C.L., C.T.M., M.C., H.B.G., Y.N., H.R. and A.E. performed the research. K.M.G. assisted in FACS experiments. T.Y.O and V.R. performed computational analysis of antibody sequences. C.O.B. performed cryo-EM data collection, processing and model building. Z.W., C.O.B., R.G., A.P.W., M.A.M., P.J.B., M.C.N. and A.E. analyzed the data. M.S.S. supervised *in vitro* neutralization assays. A.G. supervised antibody production. Z.W., C.O.B., R.G., M.A.M., P.J.B., M.C.N and A.E. wrote the manuscript.

## Acknowledgments

We thank members of the Bjorkman, Martin and Nussenzweig laboratories for discussions. Cryo-EM was performed in the Beckman Institute Resource Center for Transmission Electron Microscopy at Caltech with assistance from directors A. Malyutin and S. Chen. We thank J. Vielmetter and the Beckman Institute Protein Expression Center at Caltech for protein production, John Moore (Weill Cornell Medical College) for the BG505 stable cell line and Rogier W. Sanders (Amsterdam UMC) and Marit J. van Gils (Amsterdam UMC) for providing BG505 SOSIP trimers. This work was supported by NIH Center for HIV/AIDS Vaccine Immunology and Immunogen Discovery (CHAVI-ID) 1UM1 AI100663-01 (to M.C.N.), the National Institute of Allergy and Infectious Diseases (NIAID) HIVRAD P01 AI100148 (to M.C.N. and P.J.B.), the Bill and Melinda Gates Foundation Collaboration for AIDS Vaccine Discovery (CAVD) grant INV-002143 (to M.A.M, M.C.N, P.J.B.), the Intramural Research Program of the NIAID, NIH (to M.A.M.) and the Bill and Melinda Gates Foundation (CAVD) grant #OPP1146996 (to M.S.S). Additional support included the NIH K99/R00 grant (9694871) (to A.E.), the HHMI Hanna Gray Fellowship and the Postdoctoral Enrichment Program from the Burroughs Welcome Fund (to C.O.B.).

## Notes

### Competing Interest Statement

The authors have declared no competing interest.

