## Supplementary Materials for "A Broadly Neutralizing Macaque Monoclonal Antibody Against the HIV-1 V3-Glycan Patch"

**Supplementary Information**

**Figure S1**. **Antibodies isolated from a $SHIV_{AD8}$ infected rhesus macaque**. **A**, FACS plots show the gating strategy used to isolate single germinal center B cells that bind to YU2 gp140-F and BG505 SOSIP from a lymph node sample collected from the elite neutralizer macaque CE8J. The sorted population of B cells is highlighted in red. **B**, Pie chart shows the clonal analysis of antibodies cloned from macaque CE8J. Expanded clones are represented in colored slices while singles are shown in white. The number of analyzed sequences is shown in the middle of the pie chart. **C**, Graph shows the number of nucleotide mutations in the VH, VL and VK genes of the antibodies shown in B. **D**, Length of CDRH3 for antibodies shown in B and C. The red dot corresponds to Ab 1485. **E**, Neutralization activity of Ab1538, Ab1542 and Ab1485 determined in TZM-bl assays against a panel of 7 tier 1B and tier 2 pseudoviruses.

**Figure S2. Cryo-EM data processing and validation.** (A) Representative micrograph and 2D class averages shown for Ab1485-BG505-8ANC195 complex data. (B) Angular distribution, (C) Gold-standard Fourier shell correlation plot, with dashed lines showing resolutions at 0.5 and 0.143. (D) Local resolution as determined by ResMap depicted on side and top views of reconstructed volume. (E) Representative density contoured at 3 sigma for different regions of the complex

**Figure S3. Binding assays and structural comparison to V1//V3 antibodies raised in animals.** (A-C). SPR sensorgrams for binding of Ab1485 Fab to immobilized BG505 (A), RC1 (B), or RC1-glycan KO $^{324}$GAIA$^{327}$ (C). Sensorgrams traces are shown in colors; the fits to a 1:1

binding model are shown in black. (D) Comparison of Ab1485 (purple shades) Env binding orientation to antibodies $874_{NHP}$ (PDB: 6ORN; orange), $897_{NHP}$ (PDB: 6ORO; green), and $43A2_{rabbit}$ (PDB:6VO0; blue). Comparison with the predominant V1/V3 binding orientation observed after repeated challenge with $SHIV_{BG505}$ is also shown (modeled Fab (olive) based on cryo-EM density (EMD-20396).

**Figure S4**. **Characterization of Ab1485**. **A**, ELISA graphs showing binding of Ab1485 to lipopolysaccharide (LPS), Keyhole limpet hemocyanin (KLH), Insulin, single stranded DNA (ssDNA) and double stranded DNA (dsDNA). Antibody ED38(75, 76) and antibody mGO53(75)were used as positive and negative controls respectively. **B**, Graph shows the antibody concentrations of Ab1485-macaque-LS and the control 10-1074 in the serum of mice that carry a null mutation for the mouse neonatal Fc receptor (FcRn) and a transgene for the human FcRn at different time points after intravenous infusion of the antibodies.

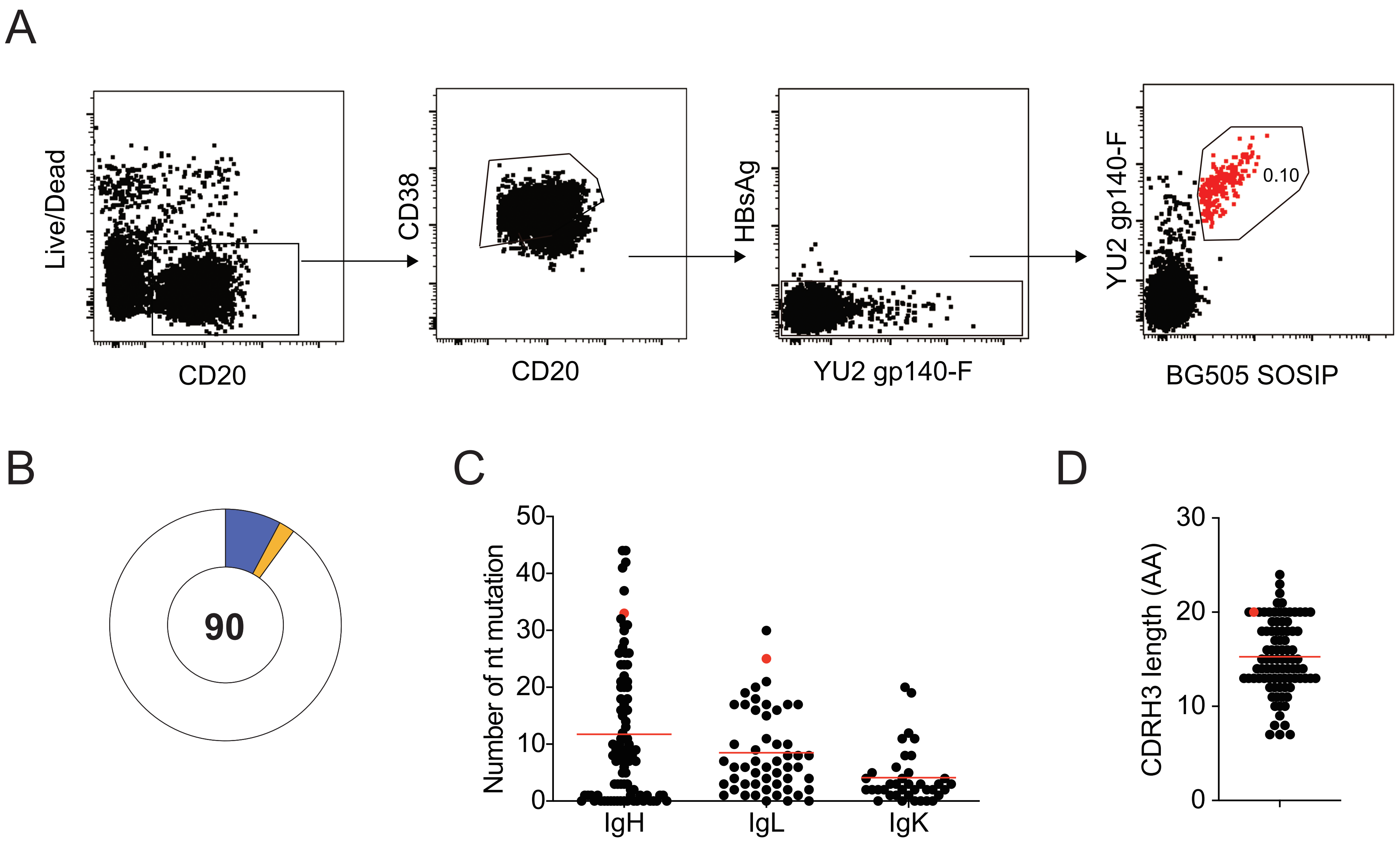


E

| Virus ID | Ab1538 | Ab1542 | Ab1485 |
|---|---|---|---|
| BG505/T332N | >267 | >412 | <0.70 |
| JRCSF.JB | >267 | <1.70 | <0.70 |
| 6535.5 | >267 | 9.21 | <0.70 |
| 92RW020.2 | >267 | >412 | <0.70 |
| Du422.1 | >267 | >412 | <0.70 |
| T250-4 | >267 | 343.2 | <0.70 |
| Q23.17 | >267 | >412 | <0.70 |

IC50 µg/ml
<1
1-100

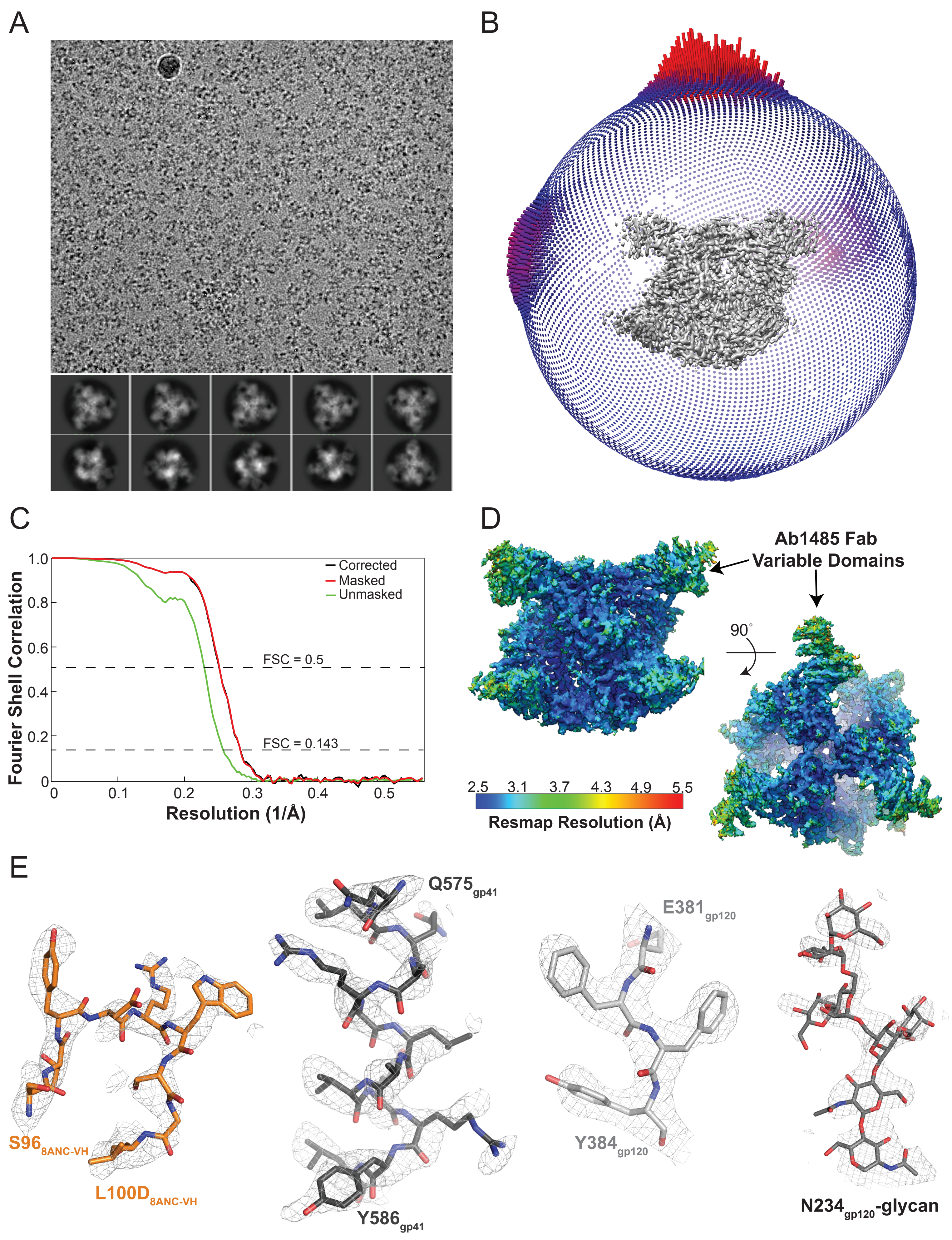

A
B
C
D
E
Fourier Shell Correlation
Resolution (1/Å)
Corrected
Masked
Unmasked
FSC = 0.5
FSC = 0.143
Ab1485 Fab
Variable Domains
90°
Resmap Resolution (Å)
2.5 3.1 3.7 4.3 4.9 5.5
S96 8ANC-VH
L100D 8ANC-VH
Q575 gp41
Y586 gp41
E381 gp120
Y384 gp120
N234 gp120-glycan

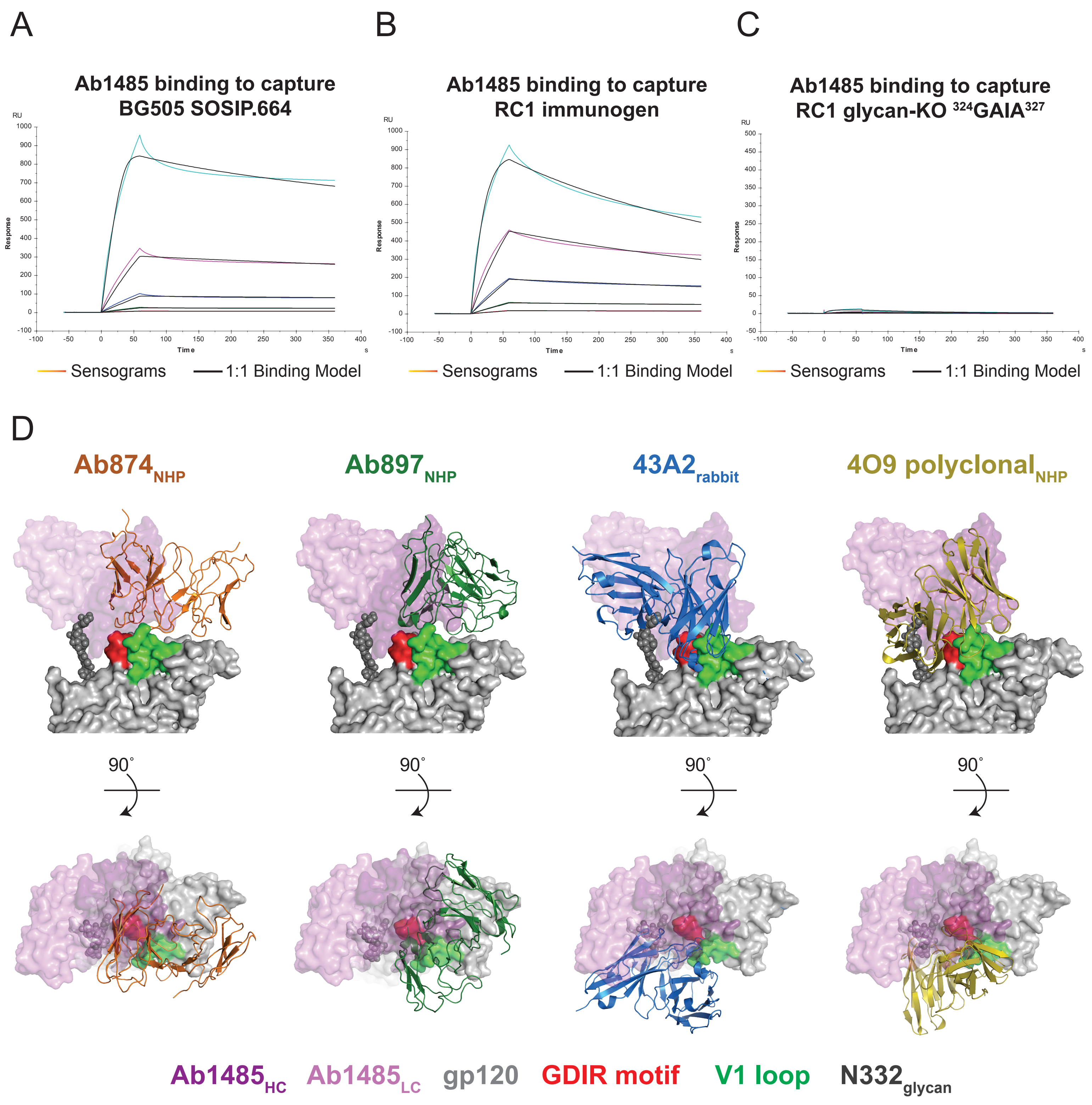

A
Ab1485 binding to capture
BG505 SOSIP.664
Response
Time
Sensograms
1:1 Binding Model
B
Ab1485 binding to capture
RC1 immunogen
Response
Time
Sensograms
1:1 Binding Model
C
Ab1485 binding to capture
RC1 glycan-KO 324GAIA327
Response
Time
Sensograms
1:1 Binding Model
D
Ab874NHP
Ab897NHP
43A2rabbit
4O9 polyclonalNHP
90°
Ab1485HC
Ab1485LC
gp120
GDIR motif
V1 loop
N332glycan

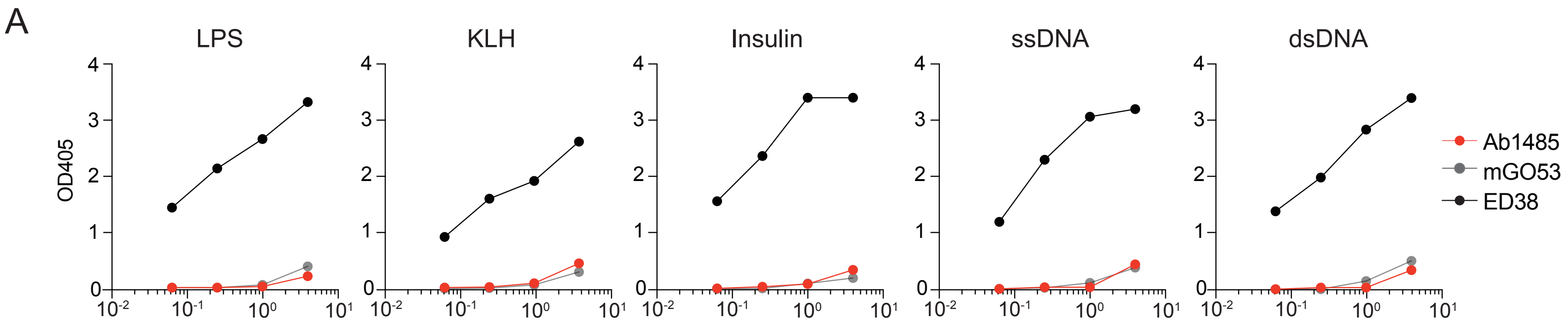
A
LPS
KLH
Insulin
ssDNA
dsDNA
OD405
Ab1485
mGO53
ED38

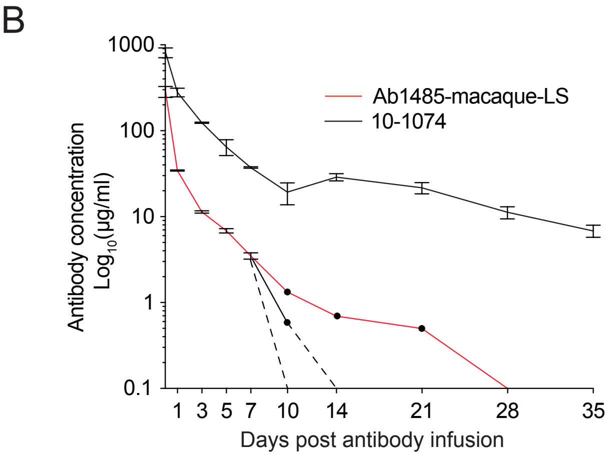
B
Ab1485-macaque-LS
10-1074
Antibody concentration
Log10(µg/ml)
Days post antibody infusion
